# Determinants of natriuretic, diuretic, and kaliuretic effects of diuretics: Sex and administration time

**DOI:** 10.1101/2022.12.03.519003

**Authors:** Pritha Dutta, Mehrshad Sadria, Anita Layton

## Abstract

Sex differences in renal function and blood pressure have been widely described across many species. Blood pressure dips during sleep and peaks in the early morning. Similarly, glomerular filtration rate, filtered electrolyte loads, urine volume, and urinary excretion all exhibit notable diurnal rhythms, which reflect, in part, the regulation of renal transporter proteins by circadian clock genes. That regulation is sexually dimorphic; as such, sex and time-of-day are not two independent regulators of kidney function and blood pressure. The objective of this study is to assess the effect of sex and administration time on the natriuretic and diuretic effects of loop, thiazide, and K^+^-sparing diuretics, which are common treatment for hypertension. Loop diuretics inhibit NKCC2 on the apical membrane of the thick ascending limb, thiazide diuretics inhibit NCC on the distal convoluted tubule, and K^+^-sparing diuretics inhibit ENaC on the connecting tubule and collecting duct. We simulate Na^+^ transporter inhibition using our sex- and time-of-day-specific computational models of mouse kidney function. Simulations results highlight significant sex and time-of day differences in drug response. Loop diuretics induce larger natriuretic and diuretic effects during the active phase. The natriuretic and diuretic effects of thiazide diuretics exhibit sex and time-of-day differences, whereas these effects of K^+^-sparing diuretics exhibit significant time-of-day difference in females only. Kaliuretic effect depends on the type of diuretics and time of administration. The present computational models can be a useful tool in chronotherapy, to tailor drug administration time to match the body’s diurnal rhythms to optimize the drug effect.

## Introduction

Physiological processes of organisms, from bacteria to plants to humans, often exhibit daily oscillations ^1^. These circadian rhythms are determined by the interactions between environmental signals known as “zeitgebers” (e.g., fluctuations in light intensity and temperature) and an internal circadian timekeeping system. In mammals, the circadian system has a complex architecture that consists of a central clock that resides in the brain’s suprachiasmatic nuclei (SCN) and peripheral clocks, located in almost all tissues and organs, that are tailored to the specific physiological functions of their respective tissues and organs ^1^. Light is the dominant zeitgeber for the SCN, whereas the peripheral clocks may preferentially respond to other zeitgebers, e.g., temperature, food intake, and physical activity. Together, the central and peripheral clocks influence circadian physiology and behavior via neuronal and humoral cues.

Diurnal variations have been observed in key components of the cardiovascular system. In particular, blood pressure fluctuates with a pattern that follows a circadian rhythm, with a peak in the early morning hours and a trough during sleep ^2^. This rhythm originates in the central clock in the SCN, but is also influenced by internal factors (e.g., kidney function, hormones, and nervous system) and external factors (e.g., light, temperature, food intake, and rest/activity routine), many of which exhibit their own rhythms. Nocturnal dipping of blood pressure is part of this normal circadian pattern, and its absence (non-dipping) has a higher prevalence among hypertensive patients, constitutes a risk factor for cardiovascular events, and is associated with more severe end-organ damage ^3^.

Besides the circadian clock, another key modulator of blood pressure regulation is sex. Indeed, sex differences in blood pressure and its dysregulation are been reported in many mammalian and avian species ^4^. In animal models of hypertension (e.g., spontaneously hypertensive rats and Dahl salt-sensitive rats), males develop earlier and more severe hypertension than females ^5^. Pre-menopausal women have lower blood pressure and lower prevalence of hypertension than age-matched men ^6^. In contrast, post-menopausal women older than 60 years have a higher prevalence of hypertension compared to men of similar ages. Blood pressure regulation is mediated by the kidney and is under the influence of multiple homeostatic and hormonal systems, including the renin-angiotensin system, the sympathetic nervous system, the oxidative stress/nitric oxide system, the endothelin system, and sex steroids. Sex differences are not limited to the sex steroids but are found in all such systems, thereby contributing to the sex differences in blood pressure and hypertension ^7^.

By managing electrolyte and fluid homeostasis, the kidney is a major regulator of blood pressure. Not surprisingly, kidney function is regulated by the circadian clock, with glomerular filtration rate (GFR), filtered electrolyte loads, urine volume, and urinary excretion exhibiting significant diurnal rhythms ^8–10^. These rhythms reflect, in part, the regulation of renal transporter proteins, including the Na^+^/H^+^ Exchanger 3 (NHE3), Na^+^-K^+^-Cl^−^ cotransporter 2 (NKCC2), Na^+^-Cl^−^ Cotransporter (NCC), Epithelial Na^+^ Channels (ENaC), pendrin, and Renal Outer-Medullary K^+^ channels (ROMK) by clock proteins brain and muscle ARNT-Like 1 (BMAL1) and Period Circadian Regulator 1 (PER1) ^11–13^. Interestingly, the regulation of renal Na^+^ transport by BMAL1 differs significantly between male and female mice ^11^. Sex has an impact on kidney functions that are regulated by, and those that are independent of, the circadian clock. The kidney mass and single-nephron GFR (SNGFR) of a male mouse are both larger than female ^14,15^. The abundance patterns of membrane transporters and channels in rodent kidneys have been reported to be sexually dimorphic ^16^. For instance, a lower NHE3 abundance has been measured in the proximal tubules of female rats and mice, which implies a lower proximal transport capacity in females compared to males. The higher fractional Na^+^ distal delivery in females is handled by the augmented transport capacity in the downstream segments ^16^. As already noted, sex and time-of-day are not two independent regulators of kidney function and blood pressure; rather, the expressions of many renal Na^+^ transporters are regulated by the circadian clock, in a sexually dimorphic manner ^11^.

A common treatment for hypertension is diuretics, which increase urine output by targeting the kidneys. The three classes of diuretic medications are loop, thiazide, and K^+^-sparing diuretics. Loop diuretics inhibit NKCC2 on the apical membrane of the thick ascending limb, thiazide diuretics inhibit NCC on the distal convoluted tubule, and K^+^-sparing diuretics inhibit ENaC on the connecting tubule and collecting duct. Given the sexual dimorphism and diurnal variations in the expression levels of these transporters, we seek to answer the questions: How do the natriuretic and diuretic effects of these diuretics differ between the sexes? And how do these effects vary during the day? To answer these questions, we simulate Na^+^ transporter inhibition using our recently published computational models of water and electrolyte transport along the nephrons of the male and female mouse kidneys that represent circadian rhythms in transporter activities ^17^.

## Methodology

We conduct simulations using our recently published sex- and time-of-day specific models of the mouse kidney function ^17^. A schematic diagram of the various cell types is given in Fig. 1. These models represent the sexual dimorphism in size, renal hemodynamics, and transporter expression patterns in mice ^16,18,19^. Female mouse proximal tubules exhibit a lower NHE3 abundance and activity. Consequently, relative to males, in females the proximal tubule reabsorbs a substantially lower fraction of filtered Na^+ 16^. The higher fractional Na^+^ distal delivery in females is handled by the augmented transport capacity in the downstream distal tubular segments. Female mice exhibit a higher abundance of NKC2C, NCC, and claudin-7 along the distal nephron segments ^16^. The models also represent the circadian regulation of GFR and transporter activities in a light-dark cycle ^8^. GFR and selected transporter activities are set to vary as sinusoidal functions of time. Model parameters that exhibit circadian rhythms are summarized in Table 1 and Fig. 2.

**Table 1.** Peak times and oscillation amplitudes of model parameters that exhibit circadian rhythms. M, male; F, female.

|  | M amplitude (%) | F amplitude (%) | Peak (ZT, 0=6am) |
| --- | --- | --- | --- |
| GFR | 12 | 12 | 16 |
| NHE3 | 10 | 10 | 9 <sup>12</sup> |
| SGLT1 | 10 | 10 | 12 <sup>13,68</sup> |
| Kir4.2 | 22 | 39 | 6 <sup>11</sup> |
| H<sup>+</sup>-K<sup>+</sup>-ATPase | 37 | 35 | 6 <sup>11</sup> |
| ROMK | 60 | 40 | 18 <sup>11</sup> |
| ENaC | 14 | 19 | 6 <sup>11</sup> |
| NKCC2 | 10 | 34 | 18 <sup>11</sup> |
| pendrin | 17 | 45 | 6 <sup>11</sup> |
| NCC | NS | 10 | 18 <sup>11</sup> |
| Claudin-8 | 50 | 50 | 2 |

**Figure 1.**
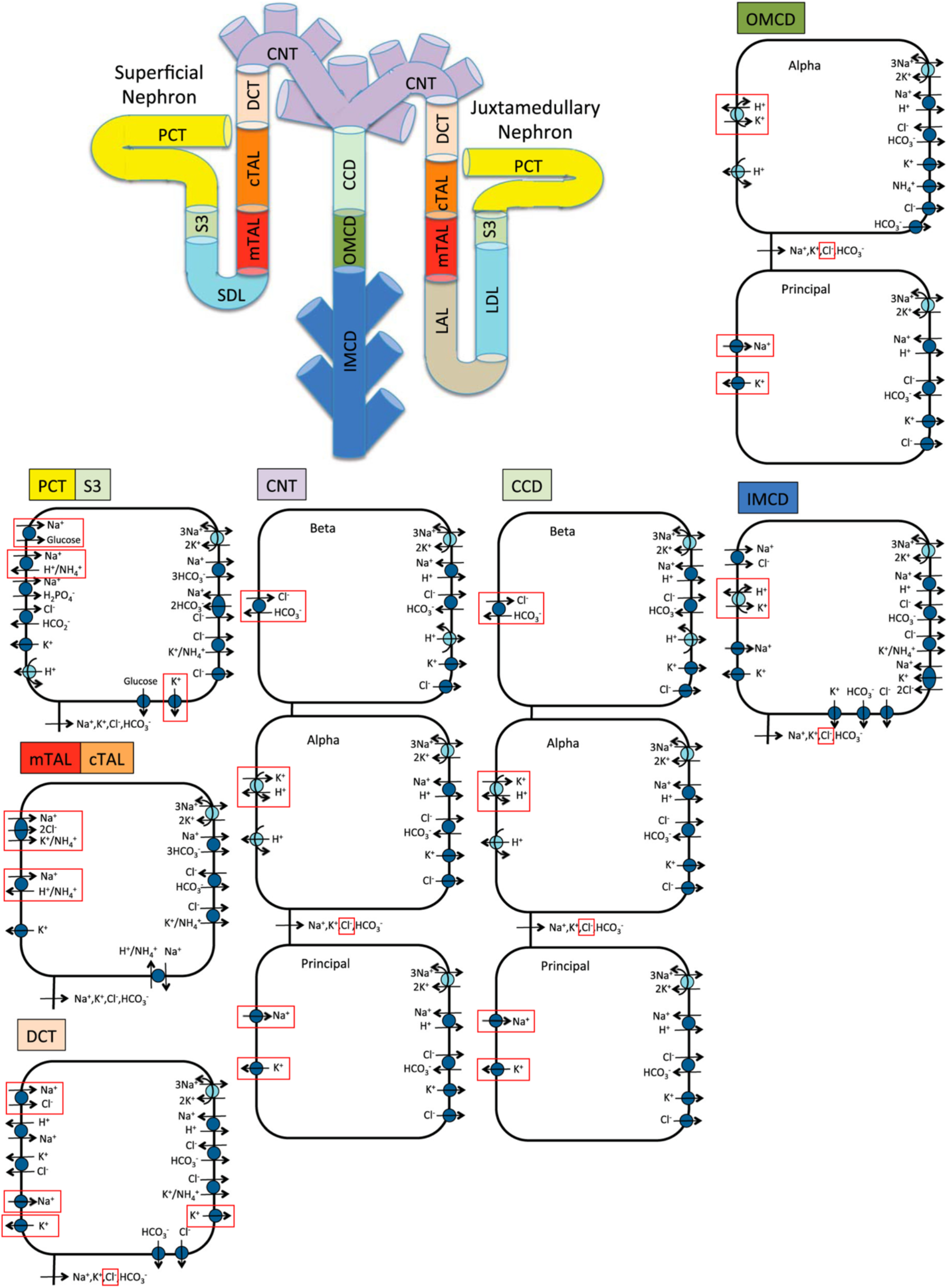
Schematic diagram of the nephron system (not to scale). The model includes one representative superficial nephron and five representative juxtamedullary nephrons, each scaled by the appropriate population ratio. Only the superficial nephron and one juxtamedullary nephron are shown. Along each nephron, the model accounts for the transport of water and 15 solutes (see text). The diagram displays only the main Na^+^, K^+^, and Cl^−^ transporters. Transporters and channels that are assumed regulated by the circadian clock are highlighted in red. In addition to solute fluxes, transmembrane and paracellular water fluxes are simulated. PCT, proximal convoluted tubule; SDL, short or outer medullary descending limb; mTAL, medullary thick ascending limb; cTAL, cortical thick ascending limb; DCT, distal convoluted tubule; CNT, connecting tubule; CCD, cortical collecting duct; OMCD, outer-medullary collecting duct; IMCD, inner medullary collecting duct; LDL, thin descending limb; LAL, thin ascending limb. Adopted from Ref. ^17^.

**Figure 2.**
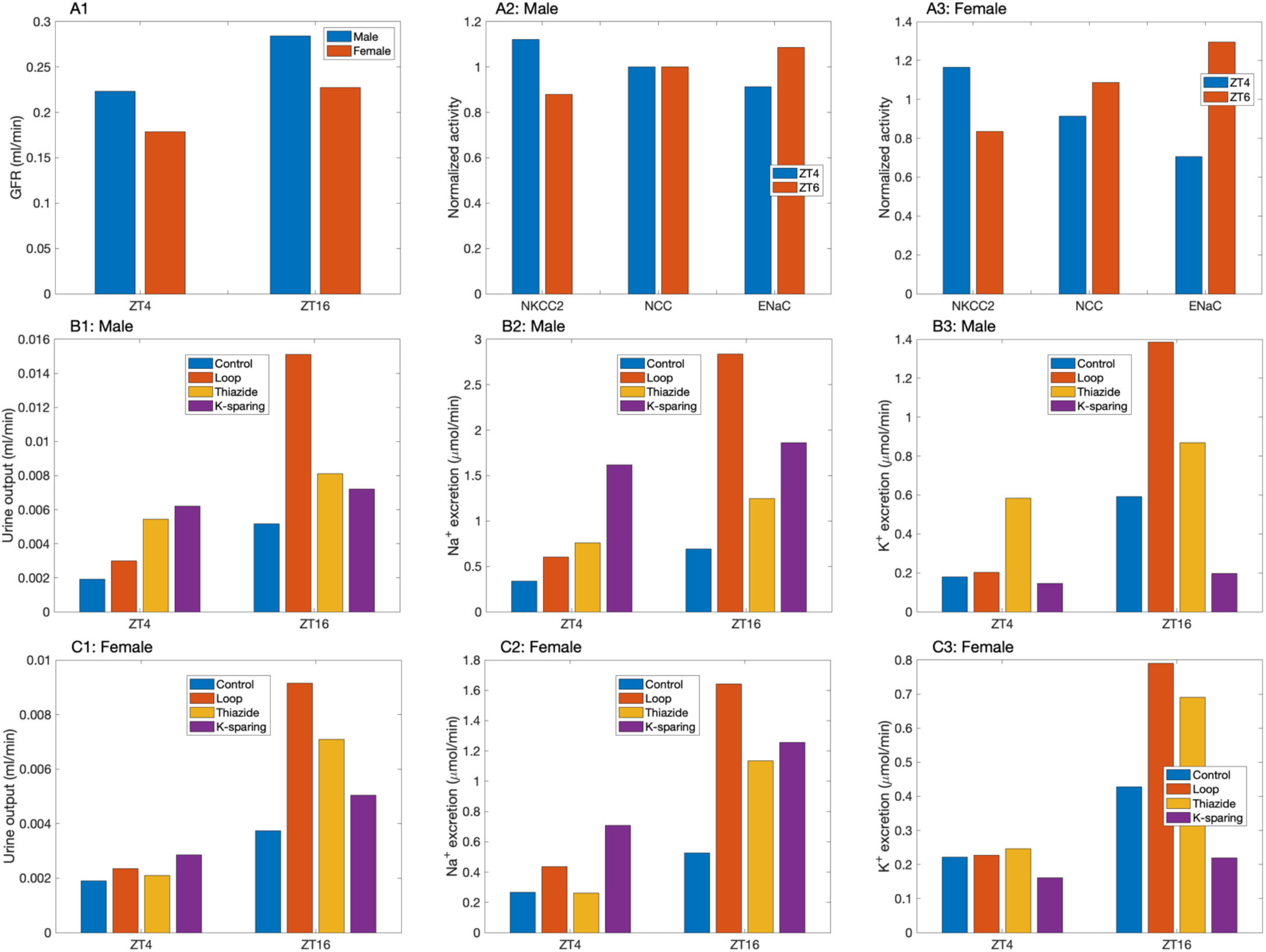
Summary of key model specifications and predictions. *A1*, GFR specified for each male and female mouse kidney, at ZT4 (light/inactive phase) and ZT16 (dark/active phase). NKCC2, NCC, and ENaC activities specified at ZT4 and ZT16, normalized by respective 24-hour mean values, for male (*A2*) and female (*A3*). Predicted urine output in male (*B1*) and female (*C1*), Na^+^ excretion (*B2, C2*), and K^+^ excretion (*B3, C3*) for control and with administration of loop, thiazide, and K^+^-sparing diuretics. In most cases, the diuretic, natriuretic, and kaliuretic effects are significantly stronger at ZT16.

### Model structure

Mouse kidneys consist of primarily superficial and juxtamedullary nephrons, with a ratio estimated to be 82:18 in male mice ^20^. In the absence of data, we assume the same ratio in females. The loops of Henle of juxtamedullary nephrons reach into differing depths of the inner medulla. To capture this heterogeneity, we represent six classes of nephrons: a superficial nephron (denoted by “SF”) and five juxtamedullary nephrons that are assumed to reach depths of 0.68, 1.36, 2.04, 2.72, and 3.4 mm (denoted by “JM-1”, “JM-2”, “JM-3”, “JM-4”, and “JM-5”, respectively) into the inner medulla. The ratios for the six nephron classes are taken to be *n*_*SF*_ = 0.82, *n*_*JM*−1_ = 0.18 × 0.4, *n*_*JM*−2_ = 0.18 × 0.3, *n*_*JM*−3_ = 0.18 × 0.15, *n*_*JM*−4_ = 0.18 × 0.1, and *n*_*JM*−5_ = 0.18 × 0.05 ^20^. The model superficial nephron includes the proximal tubule, short descending limb, thick ascending limb, distal convoluted tubule, and connecting tubule segments. Each of the model juxtamedullary nephron includes all the same segments of the superficial nephron with the addition of the long descending limbs and ascending thin limbs; these are the segments of the loops of Henle that extend into the inner medulla. The length of the long descending limbs and ascending limbs are determined by which type of juxtamedullary nephron is being modeled. The connecting tubules of the five juxtamedullary nephrons and the superficial nephron coalesce into the cortical collecting duct. SNGFR for juxtamedullary nephrons is assumed to be 20% higher than the superficial nephron SNGFR, based on superficial SNGFR and GFR measurements ^15,21^.

Each nephron segment is modeled as a tubule lined by a layer of epithelial cells in which the apical and basolateral transporters vary depending on the cell type (i.e., segment, which part of a segment, intercalated and principal cells). The models account for the following 15 solutes: Na^+^, K^+^, Cl^-^, HCO_3_^-^, H_2_CO_3_, CO_2_, NH_3_, NH_4_^+^, HPO_4_^2-^, H_2_PO_4_^-^, H^+^, HCO_2_^-^, H_2_CO_2_, urea, and glucose. The models consist of a large system of coupled ordinary differential and algebraic equations, which impose mass conservation and electroneutrality, and calculate transmembrane and paracellular fluxes ^22^. Water fluxes are driven by osmotic and hydrostatic pressure differences. Transmembrane solute fluxes may include passive and active components. An uncharged solute may be driven across a membrane passively by a concentration gradient, whereas a charged solute may be driven by an electrochemical potential gradient across an ion channel. Additional components may include coupled transport across co-transporters and/or exchangers and primary active transport across ATP-driven pumps, the activities of which may exhibit circadian rhythms ^11–13^.

Model equations are solved to predict luminal fluid flow, hydrostatic pressure, membrane potential, luminal and cytosolic solute concentrations, and transcellular and paracellular fluxes through transporters and channels. These variables are assumed to vary in space; that is, not only do they vary among different tubular segments, but these variables also vary within a specific segment, depending on the spatial location.

### Circadian rhythms in transport parameters

In a light-dark cycle, ZT0 (lights on) marks the start of the rest phase for nocturnal animals, while ZT12 (lights off) denotes the start of the active phase. The model represents the circadian rhythm in transporter activities (e.g., NHE3, NKCC2, etc., see Table 1) as

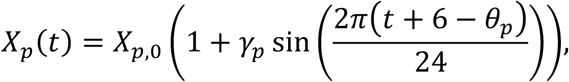

where *t* is the Zeitgeber time (ZT), *X*_*p*,0_ represents the average activity, *γ*_*p*_ denotes the fractional oscillation amplitude, and *θ*_*p*_ denotes the peak time. The circadian rhythms in GFR are represented similarly. Parameter values are specified in Table 1.

### Simulating loop diuretics

Loop diuretics inhibit NKCC2, which is expressed on the apical membrane of the thick ascending limbs of the loops of Henle. Following the procedures in Refs. ^23,24^, we assumed that the NKCC2 inhibitor was administrated for long enough to significantly impair the kidney’s ability to generate an axial osmolality gradient. Thus, we lowered the interstitial fluid concentrations of selected solutes ^23^. Given that targeted deletion of NKCC2 significantly attenuates the tubuloglomerular feedback response ^25^, we assumed that SNGFR remained at baseline values, consistent with an experimental study in the rat ^26^. We consider furosemide as a specific example of loop diuretics. Glomerular filtration of furosemide is limited; instead, it enters the proximal tubular lumen via secretion by the organic anion transporter-1 (OAT1) ^27^. The mRNA levels of OATs have been observed to exhibit circadian rhythms. For example, the expression level of OAT3 has been reported to vary by 2-folds, reaching its peak in the late light phase and early dark phase (ZT8 to ZT16) ^28^. Indeed, the diurnal expression of OATs in the kidney may contribute to time-dependent excretion of OAT substrates such as furosemide, given that deletion of a clock gene (*Bmal1*) in mice reduces the renal elimination of these substrates ^28^. We represent the effect of the diurnal expression of OATs by assuming that more NKCC2 is inhibited in the dark phase (80%) compared to the light phase (70%).

### Simulating thiazide diuretics

Thiazide diuretics inhibit NCC, which is expressed on the apical membrane of the distal convoluted tubule. In the NCC inhibition simulations, baseline interstitial concentration profiles were used. We simulate a dosage of thiazide diuretics that induces 100% inhibition of NCC. All model parameters other than NCC activities remained at baseline values.

### Simulating K^+^-sparing diuretics

K^+^-sparing diuretics inhibit ENaC, which is expressed on the apical membrane of the last third of the distal convoluted tubules as well as the full length of the connecting tubules and collecting ducts. We simulate a dosage of thiazide diuretics that induces 100% inhibition of ENaC. As in the NCC inhibition simulations, baseline interstitial concentration profiles were used and non-ENaC parameters were maintained at baseline values.

## Results

### Sex and time-of day differences in renal hemodynamics, transporter pattern, and drug response

The model represents a 24% peak-to-trough difference in GFR in both sexes, with the peak assumed at ZT16 ^9^. Activities of key transporters also exhibit circadian rhythms, including the three Na^+^ transporters analyzed in this study: NKCC2, NCC, and ENaC ^11^. Interestingly, the diurnal patterns of these three Na^+^ transporters differ. Statistically significant oscillations in NCC expression level were observed in female mice only ^11^. Expression levels of NKCC2 and NCC peak during the dark (active) phase (ZT18), whereas that of ENaC peaks half a day earlier (ZT6) ^11^ (see Figs. 1A1—1A3). The expression levels of other transporters [NHE3, Sodium-Glucose co-Transporter 1 (SGLT1), Kir4.2, H^+^-K^+^-ATPase, ROMK, pendrin, and claudin-8] vary during the day as well; see Table 1.

Changes in GFR and transporter activities together yield oscillations in urine output and excretion rates that are essentially in phase with GFR rhythms; see Figs. 2B1—2B3 and Figs. 2C1—2C3, blue bars (control). Interestingly, which drug yields the largest natriuretic and diuretic effects depends on the time of day. During the inactive phase (ZT4), K^+^-sparing diuretic has the largest effects, whereas during the active phase (ZT16), the loop diuretics dominate (Figs. 2B1, 2B2, 2C1, 2C2). Thiazide and loop diuretics produce the largest K^+^ excretion, during the inactive and active phase, respectively (Figs. 2B3 and 2C3). Qualitative similar results were predicted for males and females. Drug response is further analyzed below.

### Loop diuretics induce larger natriuretic and diuretic effects during the active phase

NKCC2 activity is assumed to peak at ZT18. Together with GFR reaching its maximum at ZT16, the model thick ascending limbs reabsorb significantly more Na^+^ at ZT16 compared to ZT4, by 18% in males and 17% in females (Fig. 3, blue bars). Administration of a loop diuretic at ZT4 is assumed to inhibit NKCC2 by 70%, resulting in 9.3% and 7.5% reduction in thick ascending limb Na^+^ reabsorption in males and females, respectively. The reduction in Na^+^ reabsorption is much less than the reduction in NKCC2 expression because the latter is partly compensated by an increase in the driving force across NKCC2 (due to a reduction in intracellular concentrations). Distal Na^+^ delivery increases, as does Na^+^ transport along the distal segments (Compare blue and red bars in Fig. 3.) Together, these processes yield similar natriuretic responses in the two sexes: a 78% increase in Na^+^ excretion in males and a 63% increase in females (Fig. 2). Changes in water transport follow Na^+^, and significant diuretic responses are predicted for males (58% increase in urine output) and females (24%).

**Figure 3.**
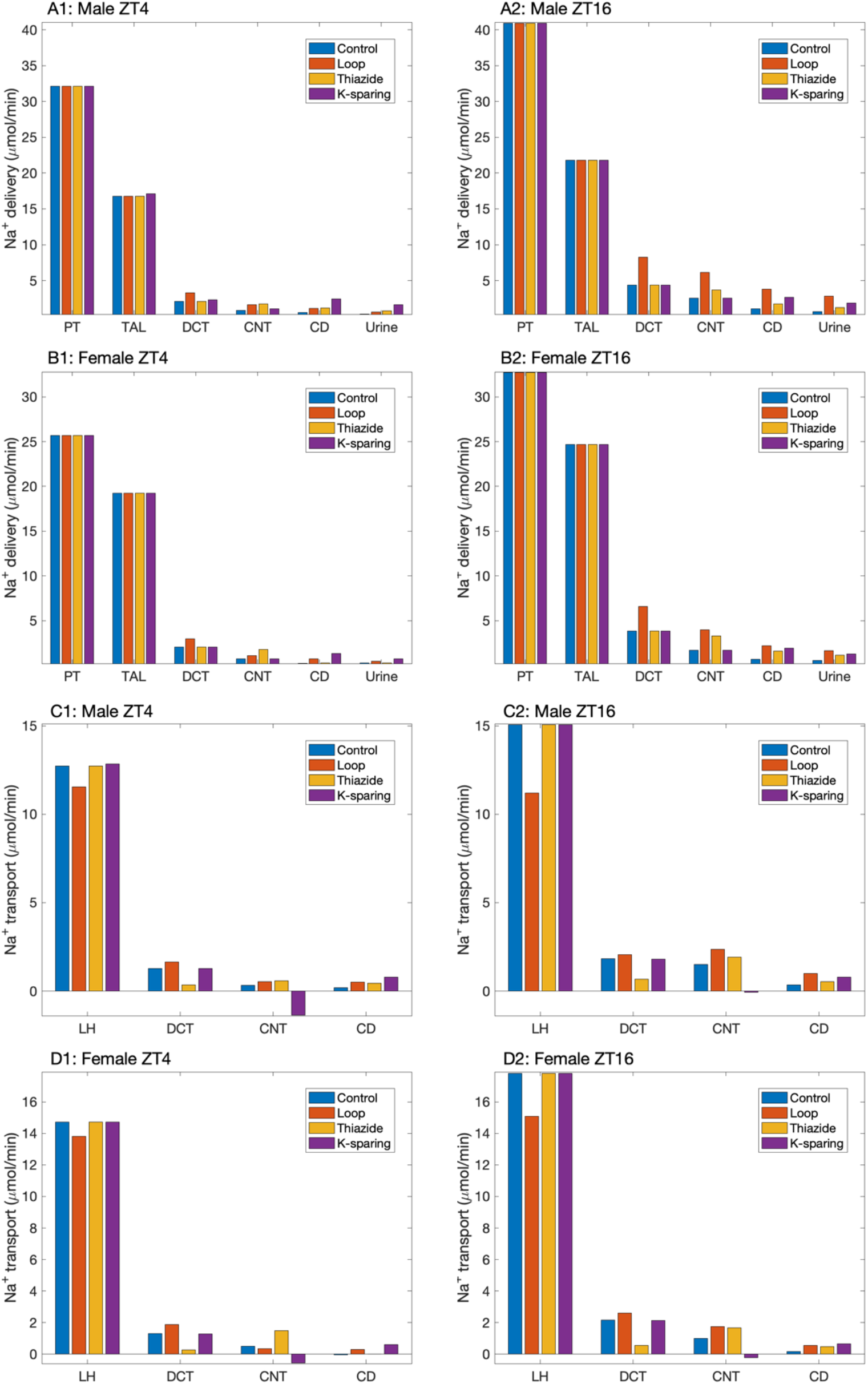
Comparison of Na^+^ delivery to individual nephron segments, given per male (*A1, A2*) and female (*B1, B2*) mouse kidney, for control and with administration of loop, thiazide, and K^+^-sparing diuretics. Results were obtained for the light/inactive phase (*A1, B1*, ZT4) and dark/active phase (*A2, B2*, ZT16) Corresponding Na^+^ transport along key segments is shown in panels *C1, C2, D1, D2*. Proximal tubule transport is not affected by these diuretics. PT, proximal tubules; LH, loops of Henle; TAL, thick ascending limbs; DCT, distal convoluted tubules; CNT, connecting tubules; CD, collecting ducts.

At ZT16, we assume that the diurnal expression of the organic anion transporters results in a stronger (80%) inhibition of NKCC2. As a result, thick ascending limb Na^+^ reabsorption decreases by 25% in males and by 16% in females. Note also that GFR and thus filtered Na^+^ is 24% higher at ZT16 than ZT4. Consequently, despite compensatory increases in Na^+^ transport along downstream nephron segments, marked natriuresis and diuresis are predicted: Na^+^ excretion increases by 316% in males and 209% in females, whereas urine output increases by 192% and 145% in males and females, respectively (see Fig. 2). Taken together, the natriuretic and diuretic responses induced by loop diuretics exhibit marked time-of-day differences ^29,30^ but, despite higher mean NKCC2 activity and NKCC2 circadian amplitude in females, only relatively limited sex differences.

### Natriuretic and diuretic effects of thiazide diuretics exhibit sex and time-of-day differences

NCC activity exhibits notable sex and time-of-day differences. Mean NCC activity is assumed to be double in females compared to males ^21,31^. Circadian rhythms were reported in females but not in males with statistical significance (Figs. 2A2 and 2A3) ^11^. These differences likely contribute to the sex and time-of-day differences in the model’s natriuretic and diuretic responses to thiazide diuretics ^32,33^.

For females, 100% inhibition of NCC substantially limits Na^+^ transport along the distal convoluted tubules. That change is partially compensated for via the upregulation of connecting tubule Na^+^ transport (Fig. 3). These competing effects result in almost no change in urine output and Na^+^ excretion at ZT4 in females (Figs. 2C1 and 2C2). In contrast, at ZT16 the circadian oscillations of GFR and NCC activity reach their peaks (Figs. 2A1 and 2A3). As such, the loss in NCC-mediated Na^+^ transport overwhelms the capacity of the downstream segments to compensate. Na^+^ transport drives water transport. Taken together, these result in 90% and 144% increases in urine output and Na^+^ excretion, respectively (Figs. 2C1 and 2C2).

For males, NCC activity is assumed to stay constant throughout the day ^11^, at a value that is half of the female’s mean ^21,31^. Because the distal Na^+^ transport capacity in males is significantly lower than females, when NCC is inhibited, downstream nephron segments fail to sufficiently compensate for the reduced Na^+^ and water transport (Fig. 3A1 and 3A2), resulting in substantial increases in urine output and Na^+^ excretion at ZT4 and ZT16 (Fig. 2).

### Natriuretic and diuretic effects of K^+^-sparing diuretics exhibit significant time-of-day difference in females only

Even though the ENaC-mediated Na^+^ reabsorption along the connecting tubules account for only a minor fraction of filtered Na^+^, ENaC inhibition gives rise to strong natriuretic and diuretic responses, in part because the collecting ducts are the only downstream segments that remain to compensate for the reduced Na^+^ transport. When ENaC is eliminated at ZT4, Na^+^ transport along the connecting tubules switches direction, from reabsorption to secretion (Figs. 3A1 and 3B1). The models predict that, among three diuretics simulated, K^+^-sparing diuretics produce the highest urine volume and Na^+^ excretion at ZT4. Note that unlike NKCC2 and NCC, whose activities peak during the active phase, ENaC activity peaks at ZT6 (inactive phase). Nonetheless, due to the higher GFR during the active phase, the connecting tubules reabsorb more Na^+^ at ZT16 than at ZT4 (Fig. 3). Hence, the administration of K^+^-sparing diuretics produces urine volume and Na^+^ excretion that are higher at ZT16 than at ZT4, although in males that difference is not as much as the analogous differences found in other diuretics (Fig. 1).

### Kaliuretic effect depends on the type of diuretics and time of administration

Loop diuretics are predicted to be K^+^ wasting but only when administered during the active phase (ZT16). At ZT4, the 70% inhibition of NKCC2 does not significantly impact K^+^ transport along the thick ascending limbs in either sex, nor does it significantly increase Na^+^ reabsorption along the connecting tubules (Figs. 5A1 and 5B1), which would have resulted in K^+^ secretion. Consequently, no significant kaliuresis is predicted to be induced by the loop diuretics at ZT4. In contrast, loop diuretics are predicted to yield the largest K^+^ excretion when administered at ZT16. The (stronger) 80% inhibition of NKCC2, together with the higher filtered K^+^, results in significantly elevated K^+^ flows into the distal convoluted tubules and connecting tubules (Fig. 4). K^+^ excretion is predicted to be 136% and 84% above baseline in male and female, respectively, the highest among all three diuretics in both sexes.

**Figure 4.**
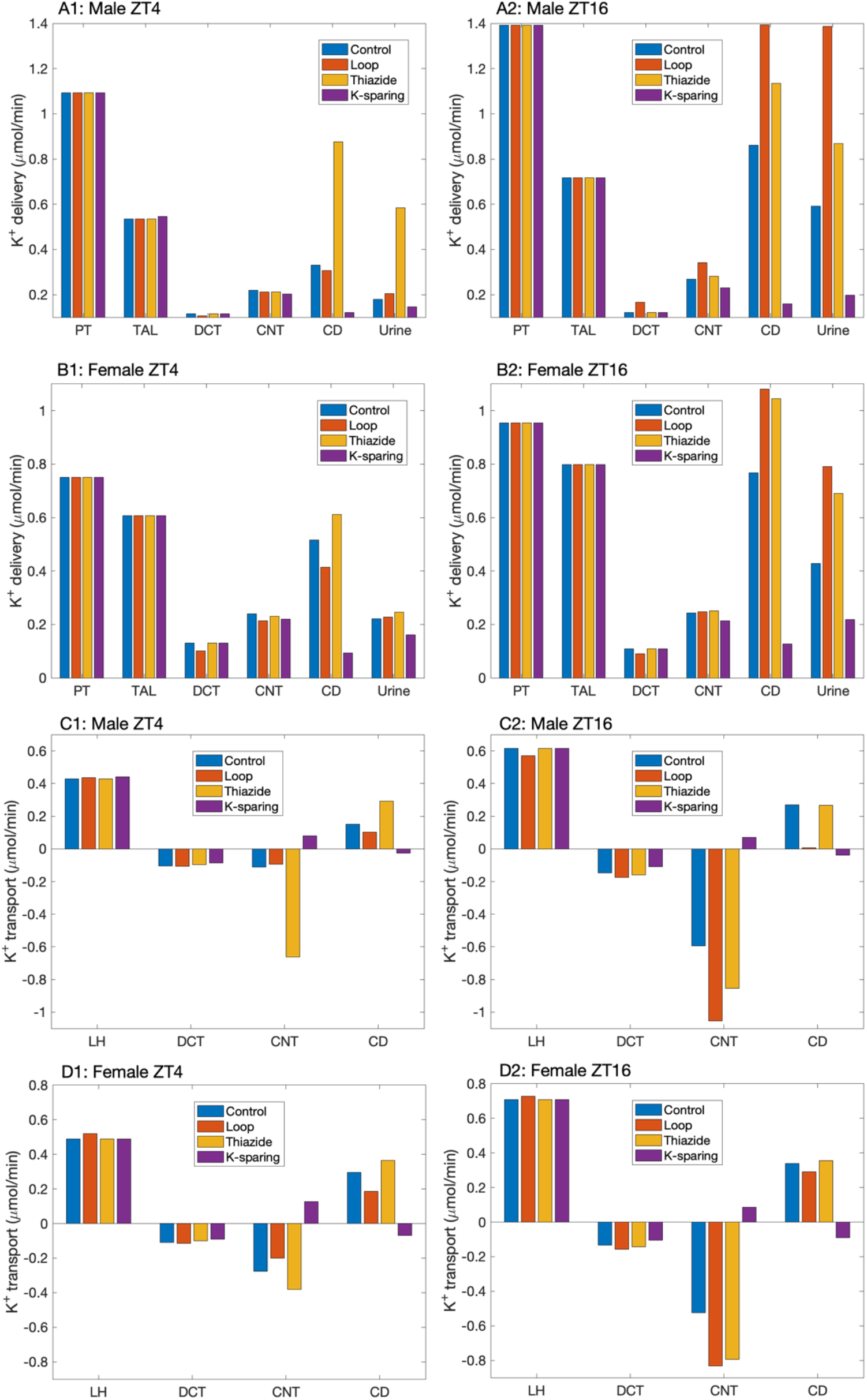
Comparison of K^+^ delivery to individual nephron segments, given per male (*A1, A2*) and female (*B1, B2*) mouse kidney, for control and with administration of loop, thiazide, and K^+^-sparing diuretics. Results were obtained for the light/inactive phase (*A1, B1*, ZT4) and dark/active phase (*A2, B2*, ZT16) Corresponding K^+^ transport along key segments is shown in panels *C1, C2, D1, D2*. Proximal tubule transport is not affected by these diuretics. PT, proximal tubules; LH, loops of Henle; TAL, thick ascending limbs; DCT, distal convoluted tubules; CNT, connecting tubules; CD, collecting ducts.

**Figure 5.**
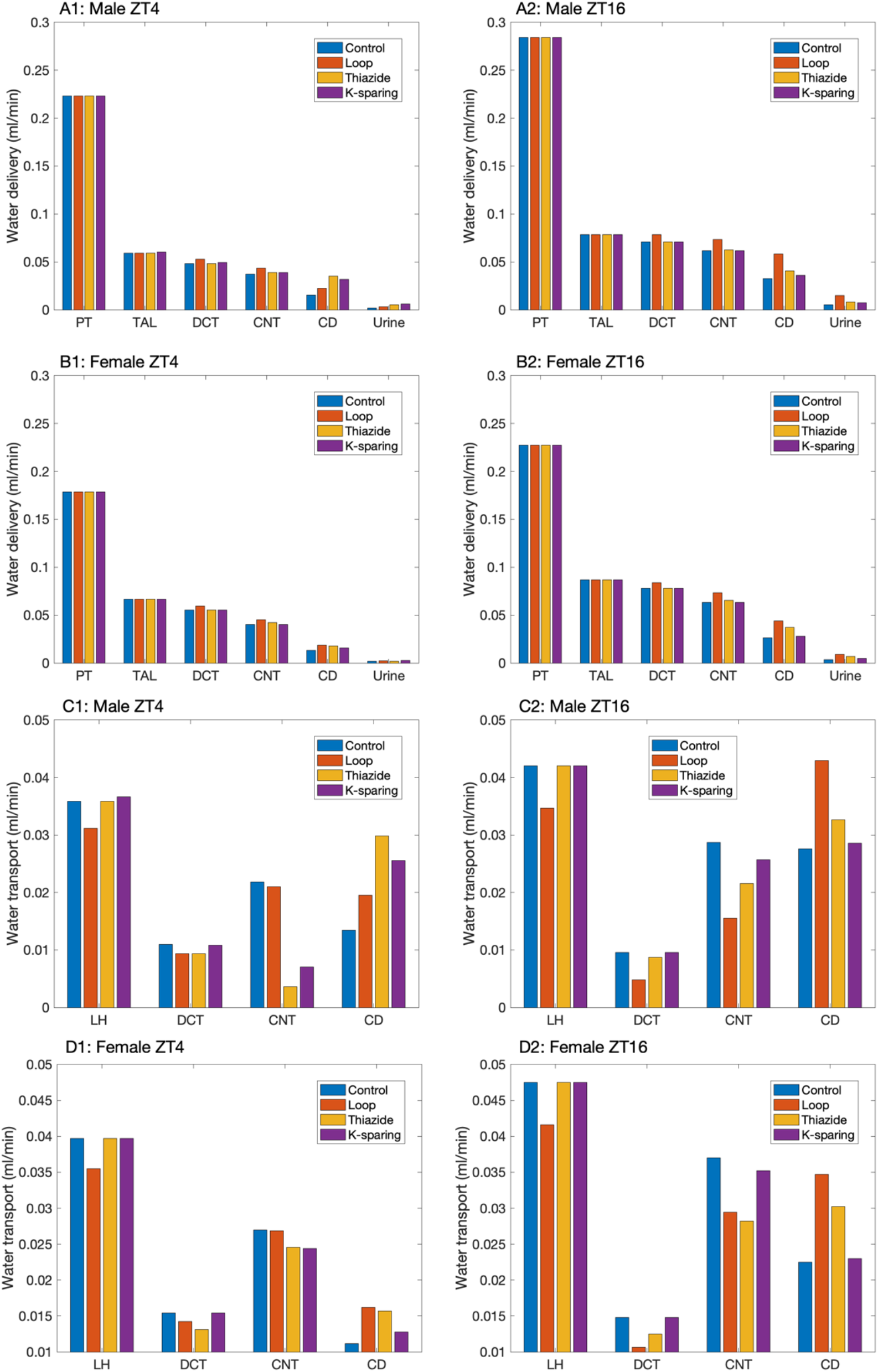
Comparison of water delivery to individual nephron segments, given per male (*A1, A2*) and female (*B1, B2*) mouse kidney, for control and with administration of loop, thiazide, and K^+^-sparing diuretics. Results were obtained for the light/inactive phase (*A1, B1*, ZT4) and dark/active phase (*A2, B2*, ZT16) Corresponding water transport along key segments is shown in panels *C1, C2, D1, D2*. Proximal tubule transport is not affected by these diuretics. PT, proximal tubules; LH, loops of Henle; TAL, thick ascending limbs; DCT, distal convoluted tubules; CNT, connecting tubules; CD, collecting ducts.

Thiazide diuretics inhibit NCC and increase Na^+^ reabsorption along the connecting tubules, thereby driving K^+^ secretion. Significant kaliuresis follows, especially during the active phase: K^+^ excretion is predicted to increase by 47% and 60% above baseline in males and females, respectively.

K^+^-sparing diuretics reduce K^+^ excretion below baseline (Figs. 2B3 and 2C3). When ENaC is inhibited, K^+^ transport switches from secretion to reabsorption, at an amount that is similarly small at ZT4 and ZT16 and in both sexes. The resulting K^+^ excretion that does not exhibit marked sex or time-of-day differences (Fig. 2).

## Discussion

Essentially every cell in our body has an internal clock. The interactions among these peripheral clocks, and with the central clock, give rise to circadian rhythms found in essentially all physiological functions. Several aspects of the cardiovascular and renal systems, e.g., blood pressure, heart rate, GFR, renal transporter expression, and urinary output, have all been reported to exhibit circadian rhythms. In particular, blood pressure follows a circadian rhythm, with values 10% to 15% lower during nighttime than during daytime (a.k.a. dipping). Concomitantly, many cardiovascular events, such as sudden cardiac death, myocardial infarction and stroke, display diurnal variations with an increased incidence in the morning ^34,35^. The absence of a nocturnal blood pressure decrease (i.e., non-dipping) is associated with target organ damage. Despite the importance and prevalence of time of day in physiological and pathophysiological processes, variations across the day are often ignored in the design and reporting of basic science and clinical research. That said, the importance of circadian rhythms in physiological function is getting increasing recognition. In recent years, there has been a growing number of experimental studies focusing on the circadian oscillations or diurnal changes in cardiovascular and kidney function (e.g., Refs. ^36–38^).

One aspect of the chronobiology of the cardiovascular system that is poorly understood is the determinants of the aforementioned dipping phenotype or its disappearance. The difficulty lies in the complexity of the regulatory networks: Multiple homeostatic and hormonal systems are involved, including the renin-angiotensin system, the sympathetic nervous system, the oxidative stress/nitric oxide system, and the endothelin system. Not only do many of these systems have competing roles, but their interactions are complex and sometimes incompletely characterized. Indeed, due to the multiple feedback loops and regulatory mechanisms, it is challenging to understand the biological consequences of diurnal or pathophysiological changes. A promising methodology for interpreting data and untangling the interactions among the multitude of regulatory processes and systems is computational modeling. Computational modeling of the circulatory system for long-term blood pressure control was pioneered by the 1972 study by Guyton et al. ^39^. A series of computational models have been developed since, e.g., Refs. ^40–44^. Surprisingly, none of these models represent the circadian rhythms in the regulatory processes of the blood pressure system. Also neglected is the effect of sex hormones, with the exceptions of sex-specific models of the renin-angiotensin system ^45^ and the blood pressure regulation system ^46^.

Equally limited are computational modeling efforts that consider the influence of sex and time of day on kidney function. Particularly relevant to blood pressure regulation are computational models of epithelial transport of electrolytes and water along the nephrons. The majority of such models were developed for the rat ^23,47–51^, or more accurately, the male rat. The marked differences in renal hemodynamics and transporter pattern ^16,18,52^ eventually inspired the development of electrolyte transport models for the female rat ^53–55^, woman ^24,56^, and female mouse ^19,57^. Similarly, computational models of kidney function have largely ignored the effect of diurnal variations, with only a few exceptions ^17,58^.

Given that intrarenal and extrarenal processes that regulate blood pressure are sex- and time-of-day-dependent, one would expect anti-hypertensive medications to exhibit sex and circadian time-dependency in their pharmacokinetics and pharmacodynamics. In this study, we focus on some aspects of one class of anti-hypertensive medications, namely, the natriuretic, diuretic, and kaliuretic effects of diuretics. Pharmacokinetics, i.e., drug absorption, distribution, metabolism, and excretion, is not considered. Urine excretion rates are predicted using a nephron model, which does not account for feedbacks via hormonal or nervous systems and thus does not fully capture sex and circadian time-dependency in pharmacodynamics. Nonetheless, a better understanding of how the mechanism of action of diuretics varies during the day and between the sexes, even in an isolated nephron system, is worthwhile. Simulations of three classes of diuretics (loop, thiazide, and K^+^-sparing) predict that the relative effect on urinary excretion depends on the time of day (Fig. 2). During the day (inactive), the K^+^-sparing diuretic has the largest natriuretic and diuretic effects, whereas during the night (active), the loop diuretics dominate. These results may be attributed to the differing circadian phases of the target transporter abundance: K^+^-sparing diuretics inhibit ENaC, which peaks during the day, whereas loop diuretics target NKCC2, which peaks at night. Loops diuretics also produces the largest K^+^ excretion when administered at night. During the inactive phase, kaliuresis is largest with thiazide diuretics (Fig. 2).

Circadian rhythms can alter the absorption, distribution, metabolism, and excretion rates of drugs. Thus, a fixed dose of drug may result in varying responses depending on the time of administration. Currently, anti-hypertensive drugs are generally prescribed for morning administration. However, several studies have shown that anti-hypertensive drugs are more effective when administered during bedtime ^59–66^. Very few medications are administered based on the time of day. A recent study of 12 different organs in a mouse model demonstrated that the protein targets of 56% of the top-selling World Health Organization approved medicines exhibit circadian oscillations ^67^. Thus, chronotherapy, i.e., tailoring drug administration time to match the body’s diurnal rhythms to increase the drug effect, can play a major role in improving modern personalized medicine. In this regard, time-of-day specific computational models would be useful tools to understand how drug responses vary across the day to determine the most effective time of administration.

## Acknowledgements

This study is supported in part by grants from the Natural Sciences and Engineering Research Council (NSERC) and Canadian Institutes of Health Research (CIHR) of Canada to A.T. Layton.

## References

1. Mohawk, J. A., Green, C. B. & Takahashi, J. S. Central and peripheral circadian clocks in mammals. Annual review of neuroscience 35, 445 (2012).

2. Biaggioni, I. Circadian Clocks, Autonomic Rhythms, and Blood Pressure Dipping. Hypertension 52, 797–798 (2008).

3. Staessen, J. A. et al. Task Force II: blood pressure measurement and cardiovascular outcome. Blood Press Monit 6, 355–370 (2001).

4. Sandberg, K. & Ji, H. Sex differences in primary hypertension. Biology of Sex Differences 3, 1–12 (2012).

5. Ouchi, Y., Share, L., Crofton, J. T., Iitake, K. & Brooks, D. P. Sex difference in the development of deoxycorticosterone-salt hypertension in the rat. Hypertension 9, 172–177 (1987).

6. Gu, Q., Burt, V. L., Paulose-Ram, R. & Dillon, C. F. Gender Differences in Hypertension Treatment, Drug Utilization Patterns, and Blood Pressure Control Among US Adults With Hypertension: Data From the National Health and Nutrition Examination Survey 1999–2004. American Journal of Hypertension 21, 789–798 (2008).

7. Reckelhoff, J. F. Sex Differences in Regulation of Blood Pressure. in Sex-Specific Analysis of Cardiovascular Function (eds. Kerkhof, P. L. M. & Miller, V. M.) 139–151 (Springer International Publishing, 2018). doi:10.1007/978-3-319-77932-4_9.

8. Johnston, J. G. & Pollock, D. M. Circadian regulation of renal function. Free Radical Biology and Medicine 119, 93–107 (2018).

9. Pons, M., Tranchot, J., L’azou, B. & Cambar, J. Circadian rhythms of renal hemodynamics in unanesthetized, unrestrained rats. Chronobiology international 11, 301–308 (1994).

10. Poulis, J., Roelfsema, F. & Van Der Heide, D. Circadian urinary excretion rhythms in adrenalectomized rats. American Journal of Physiology-Regulatory, Integrative and Comparative Physiology 251, R441–R449 (1986).

11. Crislip, G. R. et al. Differences in renal BMAL1 contribution to Na+ homeostasis and blood pressure control in male and female mice. American Journal of Physiology-Renal Physiology 318, F1463–F1477 (2020).

12. Rohman, M. S. et al. Circadian clock genes directly regulate expression of the Na+/H+ exchanger NHE3 in the kidney. Kidney international 67, 1410–1419 (2005).

13. Solocinski, K. et al. Transcriptional regulation of NHE3 and SGLT1 by the circadian clock protein Per1 in proximal tubule cells. American Journal of Physiology-Renal Physiology 309, F933–F942 (2015).

14. Laouari, D. et al. The sexual dimorphism of kidney growth in mice and humans. Kidney International (2022).

15. Messow, C., Gärtner, K., Hackbarth, H., Kangaloo, M. & Lünebrink, L. Sex differences in kidney morphology and glomerular filtration rate in mice. Contributions to Nephrology 19, 51–55 (1980).

16. Veiras, L. C. et al. Sexual Dimorphic Pattern of Renal Transporters and Electrolyte Homeostasis. J Am Soc Nephrol 28, 3504–3517 (2017).

17. Layton, A. T. & Gumz, M. L. Sex Differences in Circadian Regulation of Kidney Function of the Mouse. American Journal of Physiology-Renal Physiology (2022) doi:10.1152/ajprenal.00227.2022.

18. Munger, K. & Baylis, C. Sex differences in renal hemodynamics in rats. Am J Physiol 254, F223–31 (1988).

19. Stadt, M. M. & Layton, A. T. Sex and Species Differences in Epithelial Transport in the Rat and Mouse Kidneys: Modeling and Analysis. Frontiers in physiology (2022).

20. Zhai, X.-Y. et al. Three-dimensional reconstruction of the mouse nephron. Journal of the American Society of Nephrology 17, 77–88 (2006).

21. Xu, S. et al. Sex difference in kidney electrolyte transport III: Impact of low K intake on thiazide-sensitive cation excretion in male and female mice. Pflügers Archiv-European Journal of Physiology 473, 1749–1760 (2021).

22. Layton, A. A Complete Set of Equations for a Computational Model of Electrolyte and Water Transport along the Nephrons in a Mammalian Kidney. 2022.09.23.509286 Preprint at https://doi.org/10.1101/2022.09.23.509286 (2022).

23. Layton, A. T., Laghmani, K., Vallon, V. & Edwards, A. Solute transport and oxygen consumption along the nephrons: effects of Na+ transport inhibitors. Am J Physiol Renal Physiol 311, F1217–F1229 (2016).

24. Hu, R., McDonough, A. A. & Layton, A. T. Sex differences in solute and water handling in the human kidney: Modeling and functional implications. PLoS Comput Biol (2021).

25. Johnston, P. A. & Kau, S. T. The effect of loop of Henle diuretics on the tubuloglomerular feedback mechanism. Methods Find Exp Clin Pharmacol 14, 523–9 (1992).

26. Romano, G., Favret, G., Federico, E. & Bartoli, E. The site of action of furosemide. Pharmacol Res 37, 409–19 (1998).

27. Wilcox, C. S. New Insights into Diuretic Use in Patients with Chronic Renal Disease. JASN 13, 798–805 (2002).

28. Nikolaeva, S. et al. Nephron-Specific Deletion of Circadian Clock Gene Bmal1 Alters the Plasma and Renal Metabolome and Impairs Drug Disposition. JASN 27, 2997–3004 (2016).

29. Renal Adaptations to Continuous Administration of Furosemide and Bendroflumethiazide in Rats - Lunau - 1994 - Pharmacology & Toxicology - Wiley Online Library. https://onlinelibrary.wiley.com/doi/abs/10.1111/j.1600-0773.1994.tb01101.x?casa_token=STGXlValy6cAAAAA:Zr-jEdnGhhnH7wPChvZHB2uqMlkOaf-KSyfJWYPsiV0oEPP4rRlUXY0WhgRS4PSApwNgU02ti2zVNj.

30. Shalmi, M., Rasmusen, H., Amtorp, O. & Christensen, S. Effect of chronic oral furosemide administration on the 24-hour cycle of lithium clearance and electrolyte excretion in humans. Eur J Clin Pharmacol 38, 275–280 (1990).

31. Li, J. et al. Gender difference in kidney electrolyte transport. I. Role of AT1a receptor in thiazide-sensitive Na+-Cl− cotransporter activity and expression in male and female mice. American Journal of Physiology-Renal Physiology 313, F505–F513 (2017).

32. Fujimura, A., Ohira, H., Shiga, T., Ohashi, K. & Ebihara, A. Chronopharmacology of Trichlormethiazide in Rats. The Japanese Journal of Pharmacology 55, 294–298 (1991).

33. Circadian exosomal expression of renal thiazide-sensitive NaCl cotransporter (NCC) and prostasin in healthy individuals - Castagna - 2015 - PROTEOMICS – Clinical Applications - Wiley Online Library. https://onlinelibrary.wiley.com/doi/full/10.1002/prca.201400198?casa_token=zmVwvWyKgEwAAAAA%3AxuJC475dDv7GvkmWINs0XIfuWuGiFoyiPVWRD30Aj0AQWB-D8L_OAnUydUeh0czGnJQZHILS9oqXrg.

34. Muller, J. E. et al. Circadian variation in the frequency of onset of acute myocardial infarction. New England Journal of Medicine 313, 1315–1322 (1985).

35. Elliott, W. J. Circadian Variation in the Timing of Stroke Onset. Stroke 29, 992–996 (1998).

36. Firsov, D. & Bonny, O. Circadian rhythms and the kidney. Nature Reviews Nephrology 14, 626–635 (2018).

37. Costello, H. M., Johnston, J. G., Juffre, A., Crislip, G. R. & Gumz, M. L. Circadian Clocks of the Kidney: Function, Mechanism, and Regulation. Physiological Reviews (2022).

38. Chen, L. & Yang, G. Recent advances in circadian rhythms in cardiovascular system. Frontiers in Pharmacology 6, (2015).

39. Guyton, A., Coleman, T. & Granger, H. Circulation: Overall regulation. Annu Rev Physiol 34, 13–46 (1972).

40. Beard, D. A., Pettersen, K. H., Carlson, B. E., Omholt, S. W. & Bugenhagen, S. M. A computational analysis of the long-term regulation of arterial pressure [version 1; referees: 1 approved, 2 approved with reservations]. vol. 2 (2013).

41. Thomas, S. R. et al. SAPHIR: a physiome core model of body fluid homeostasis and blood pressure regulation. Philos Trans A Math Phys Eng Sci 366, 3175–97 (2008).

42. Hallow, K. M. et al. A model-based approach to investigating the pathophysiological mechanisms of hypertension and response to antihypertensive therapies: extending the Guyton model. American Journal of Physiology - Regulatory, Integrative and Comparative Physiology 306, R647 (2014).

43. Karaaslan, F., Denizhan, Y., Kayserilioglu, A. & Gulcur, H. O. Long-term mathematical model involving renal sympathetic nerve activity, arterial pressure, and sodium excretion. Ann Biomed Eng 33, 1607–30 (2005).

44. Averina, V. A., Othmer, H. G., Fink, G. D. & Osborn, J. W. A new conceptual paradigm for the haemodynamics of salt-sensitive hypertension: a mathematical modelling approach. The Journal of Physiology 590, 5975–5992 (2012).

45. Leete, J. G. S; Layton, AT. Modeling Sex Differences in the Renin Angiotensin System and the Efficacy of Antihypertensive Therapies. Computers & Chemical Engineering 112, 253– 264 (2018).

46. Leete, J. & Layton, A. T. Sex-specific long-term blood pressure regulation: Modeling and analysis. Comput Biol Med 104, 139–148 (2019).

47. Weinstein, A. M. A mathematical model of the rat nephron: glucose transport. Am J Physiol Renal Physiol 308, F1098–118 (2015).

48. Weinstein, A. M. A mathematical model of the rat kidney: K(+)-induced natriuresis. Am J Physiol Renal Physiol 312, F925–F950 (2017).

49. Layton, A. T. & Vallon, V. SGLT2 inhibition in a kidney with reduced nephron number: modeling and analysis of solute transport and metabolism. American Journal of Physiology-Renal Physiology 314, F969–F984 (2018).

50. Layton, A. T., Edwards, A. & Vallon, V. Renal potassium handling in rats with subtotal nephrectomy: Modeling and Analysis. Am J Physiol Renal Physiol (2017).

51. Layton, A. T., Vallon, V. & Edwards, A. A computational model for simulating solute transport and oxygen consumption along the nephrons. American Journal of Physiology-Renal Physiology 311, F1378–F1390 (2016).

52. Sgouralis, I., Evans, R. G. & Layton, A. T. Renal medullary and urinary oxygen tension during cardiopulmonary bypass in the rat. Math Med Biol 34, 313–333 (2016).

53. Hu, R., McDonough, A. A. & Layton, A. T. Sex-Differences in Solute Transport Along the Nephrons: Effects of Na+ Transport Inhibition. American Journal of Physiology-Renal Physiology (2020).

54. Hu, R., McDonough, A. A. & Layton, A. T. Functional implications of the sex differences in transporter abundance along the rat nephron: modeling and analysis. Am J Physiol Renal Physiol 317, F1462–F1474 (2019).

55. Li, Q., McDonough, A. A., Layton, H. E. & Layton, A. T. Functional implications of sexual dimorphism of transporter patterns along the rat proximal tubule: modeling and analysis. Am J Physiol Renal Physiol 315, F692–F700 (2018).

56. Swapnasrita, S., Carlier, A. & Layton, A. T. Sex-specific computational models of kidney function in patients with diabetes. Frontiers in physiology 14 (2022).

57. Fattah, H., Layton, A. & Vallon, V. How Do Kidneys Adapt to a Deficit or Loss in Nephron Number? Physiology 34, 189–197 (2019).

58. Wei, N., Gumz, M. L. & Layton, A. T. Predicted effect of circadian clock modulation of NHE3 of a proximal tubule cell on sodium transport. American Journal of Physiology-Renal Physiology 315, F665–F676 (2018).

59. Hermida, R. C. et al. Bedtime hypertension treatment improves cardiovascular risk reduction: the Hygia Chronotherapy Trial. European Heart Journal 41, 4565–4576 (2020).

60. Diuretic drugs benefit patients with hypertension more with night-time dosing - Basil N Okeahialam, Esther N Ohihoin, Jayne NA Ajuluchukwu, 2012. https://journals.sagepub.com/doi/full/10.1177/2042098612459537.

61. Hermida, R. C., Ayala, D. E., Mojón, A., Fontao, M. J. & Fernández, J. R. Chronotherapy With Valsartan/Hydrochlorothiazide Combination in Essential Hypertension: Improved Sleep-Time Blood Pressure Control With Bedtime Dosing. Chronobiology International 28, 601– 610 (2011).

62. Calvo, C. et al. Chronotherapy with torasemide in hypertensive patients: increased efficacy and therapeutic coverage with bedtime administration. Med Clin (Barc) 127, 721–729 (2006).

63. Chronotherapy in Nigerian hypertensives - Basil Okeahialam, Esther Ohihoin, Jayne Ajuluchukwu, 2011. https://journals.sagepub.com/doi/full/10.1177/1753944711402119.

64. Hermida, R. C. et al. Comparison of the Effects on Ambulatory Blood Pressure of Awakening versus Bedtime Administration of Torasemide in Essential Hypertension. Chronobiology International 25, 950–970 (2008).

65. Evening versus morning dosing regimen drug therapy for chronic kidney disease patients with hypertension in blood pressure patterns: a systematic review and meta-analysis - Wang - 2017 - Internal Medicine Journal - Wiley Online Library. https://onlinelibrary.wiley.com/doi/full/10.1111/imj.13490?casa_token=PqbDqpHCn-QAAAAA%3AnPoLhbhG93UwdB2FCqEfVuZYl4H3rdwCyJnN_WrdyB4v7MQ2lFTx9EgxFz7E54rKgEggHuD8Wfj02Q.

66. Hermida, R. C. & Smolensky, M. H. Chronotherapy of hypertension. Current Opinion in Nephrology and Hypertension 13, 501–505 (2004).

67. Zhang, R., Lahens, N. F., Ballance, H. I., Hughes, M. E. & Hogenesch, J. B. A circadian gene expression atlas in mammals: Implications for biology and medicine. Proceedings of the National Academy of Sciences 111, 16219–16224 (2014).

68. Solocinski, K. & Gumz, M. L. The circadian clock in the regulation of renal rhythms. Journal of biological rhythms 30, 470–486 (2015).

